# Structural basis for silicon uptake by higher plants

**DOI:** 10.1101/2021.04.14.439814

**Authors:** Bert van den Berg, Conrado Pedebos, Jani R. Bolla, Carol V. Robinson, Arnaud Baslé, Syma Khalid

## Abstract

Metalloids are elements with physical and chemical properties that are intermediate between metals and non-metals. Silicon (Si) is the most abundant metalloid in the Earth’s crust and occurs at high levels in many plants, especially those belonging to the Poaceae (grasses). Most of the world’s staple food crops such as rice, barley and maize accumulate silicon to high levels, resulting in resistance to abiotic and biotic stresses and consequently better plant growth and crop yields. The first step in silicon accumulation is the uptake of silicic acid (Si), the bioavailable from of silicon, by the roots, a process mediated by the structurally uncharacterised NIP subfamily of aquaporins. Here we present the X-ray crystal structure of the archetypal NIP family member from *Oryza sativa* (OsNIP2;1). While the OsNIP2;1 channel is closed in the crystal by intracellular loop D, unbiased molecular dynamics (MD) simulations reveal a rapid channel opening on sub-microsecond time scales. MD simulations further show how Si interacts with an extracellular five-residue selectivity filter that provides the main barrier for transmembrane diffusion. Our data provide a foundation for understanding and potential manipulation of metalloid selectivity of an important and understudied aquaporin subfamily.

**Significance:** Many of the world’s most important food crops such as rice, barley and maize accumulate silicon to high levels, resulting in better plant growth and crop yields. The first step in silicon accumulation is the uptake of silicic acid (Si) by the roots, a process mediated by the structurally uncharacterised NIP subfamily of aquaporins. Here, we present the X-ray crystal structure and molecular dynamics simulations of the archetypal NIP family member from *Oryza sativa* (OsNIP2;1) to visualise Si uptake. Our data provide a platform for improved understanding of Si uptake by plants that could be utilised, *e.g*., in silicon biofortification of important crops and potential alleviation of arsenic accumulation in the rice grain.

## Introduction

Many higher plants, and members of the Poaceae (grasses) in particular, accumulate silicon to high levels (up to 10% w/w for rice) (1). Silicon is generally not yet considered an essential plant element, but a high silicon content provides resistance to abiotic and biotic stresses, improves the light-interception ability by plants in a community, and minimises transpiration losses (1,2). Silicic acid (Si; H_4_SiO_4_; pK_a_ = 9.25) is the naturally occurring bioavailable form of silicon. At pH values in most soils, it is a polar but neutral molecule and soluble to concentrations of ~2 mM (3). In shoots and leaves, accumulating Si is spontaneously transformed to solid amorphous silica (SiO_2_–nH_2_O) called silica bodies, which are deposited mainly in the cell walls of different tissues and generate structural and mechanical stability (1). The facilitated diffusion of silicon and other metalloids such as boron and arsenic across bilayers is mediated by members of the NIP (<u>n</u>odulin26 intrinsic <u>p</u>rotein) subfamily of aquaporins (4), also termed metalloid porins (1). NIPs occur not only in roots but in most plant tissues, and can be divided into three functional groups, NIP-I, NIP-II and NIP-III, based on the composition of the four-residue selectivity filter (SF) or aromatic-arginine (ar/R) region at the extracellular mouth of the channel (1). NIP-I members have the most stringent substrate selectivity and appear to only transport water, glycerol and arsenous acid (H_3_AsO_3_; As). NIP-II members additionally transport boric acid (B), while members of the NIP-III subgroup have the broadest selectivity and also transport Si. Based on the predicted predominance of small residues at the SF (typically GSGR), it has been proposed that NIP-III channels may have the widest SF (1), which would explain why they transport the widest range of substrates.

*Lsi1* (<u>l</u>ow <u>si</u>licon rice 1), caused by the absence of *Oryza sativa* NIP2;1 (OsNIP2;1), is a rice mutant defective in Si uptake with various pest-sensitive phenotypes, and has a grain yield of only 10% compared to wild type rice (5). OsNIP2;1 has been characterised extensively using *Xenopus* oocytes and was found to transport Si efficiently, but B and glycerol very poorly, indicating substrate selectivity (6). OsNIP2;1 is highly expressed on the distal side of plasma membranes of root cells, and functions together with the active silicon efflux transporter Lsi2, localised in the proximal membranes of root cells, to drive unidirectional Si transport towards the xylem (7,8). Phylogenetically, NIP family members cluster together with bacterial and archaeal AqpN proteins in arsenic resistance (*ars*) operons, suggesting NIP channels may have evolved from As efflux proteins (9). Indeed, OsNIP2;1 also transports As (arsenous acid/arsenite) efficiently (10), making it a major cause of toxic arsenic accumulation in the rice grain.

## Results and Discussion

In contrast to classical aquaporins (AQPS) and aquaglyceroporins (AQGPs) (4,11), no structural information is yet available for any NIP family member, and the basis for metalloid selectivity remains therefore unclear. To address this knowledge gap, we expressed *osnip2;1* to high levels (~1 mg/liter) in *Saccharomyces cerevisiae*, followed by purification and crystallisation in detergent. Well-diffracting crystals could only be obtained from a truncated form of OsNIP2;1 generated via limited trypsinolysis, which was shown to comprise residues 38-264 by proteomics analysis. The truncated form is a tetramer in solution, as confirmed by native mass spectrometry (Fig. 1A and Fig. S1). The X-ray crystal structure was determined using data to 2.6 Å resolution, via molecular replacement with archaeal AqpM (Table S1). A low-resolution structure (3.8 Å) was also obtained for the full-length (FL) protein, and is identical to that of the truncated protein, suggesting the poorly conserved N- and C-terminal ~35 residues that are invisible in the FL structure are likely disordered. As expected from sequence similarity (Fig. S2), OsNIP2;1 displays the typical aquaporin fold, with a tetrameric assembly in which each protomer has six transmembrane helices and two half-helices in cytoplasmic loop B and extracellular loop E, which together form a seventh pseudo-transmembrane segment with the important NPA motifs in the centre of the bilayer (Fig. 1B).

**Figure 1.**
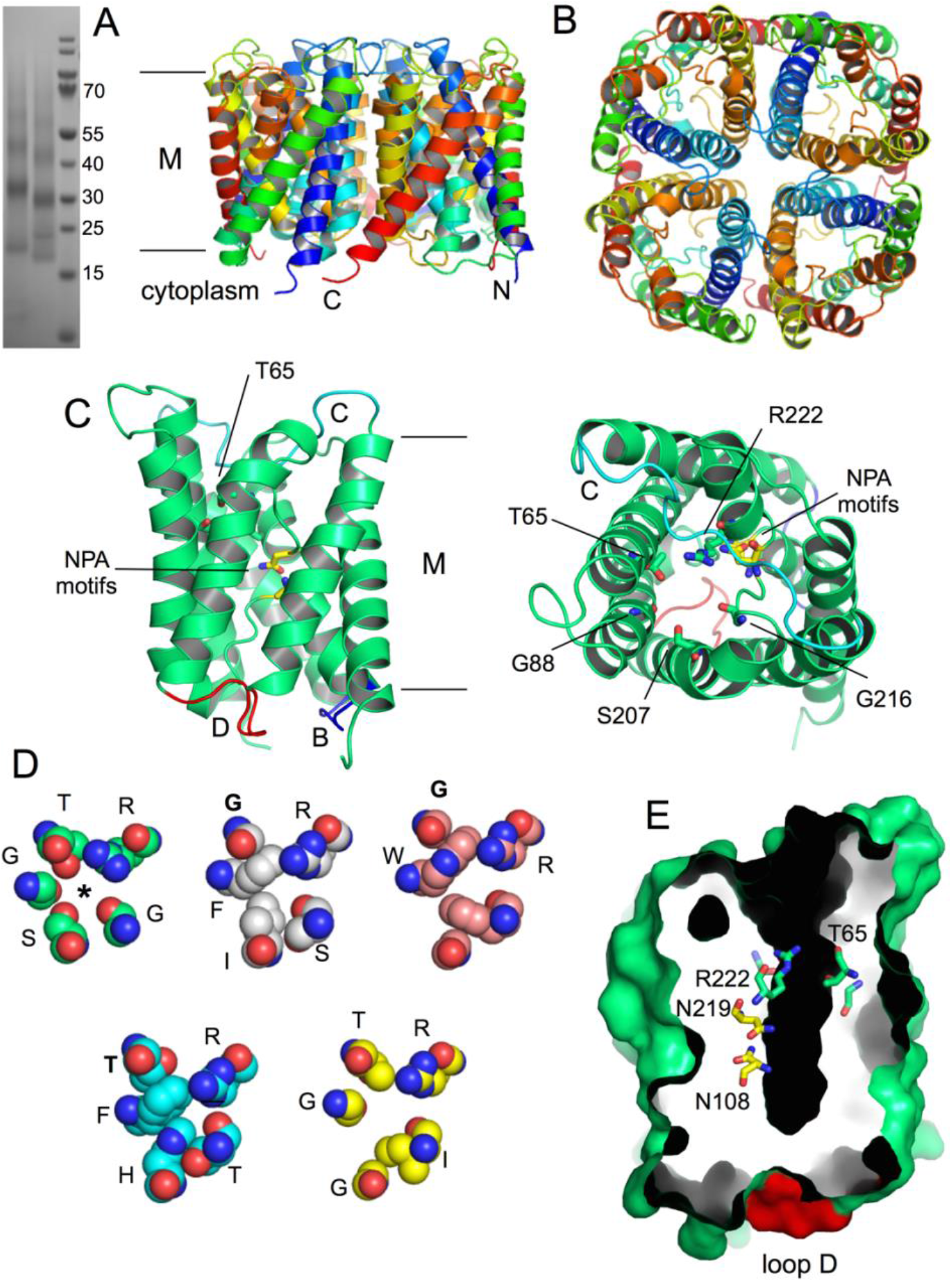
X-ray crystal structure of OsNIP2;1. (A) SDS-PAGE gel of purified OsNIP2;1 before (left lane) and after (middle lane) limited trypsinolysis. Molecular markers have been loaded in the right lane. (B) Cartoon overview of the OsNIP2;1 tetramer, rainbow coloured by chain from the N-terminus (blue) to the C-terminus (red). The approximate membrane (M) boundaries are shown. (C) Monomer cartoon viewed from the membrane plane (left panel) and from the extracellular side. Selectivity filter (SF) residues and the asparagine residues of the two NPA motifs are shown as sticks. Cytoplasmic loops B and D are coloured dark blue and red, respectively. Extracellular loop C is coloured cyan. (D) Comparison of selectivity filters of aquaporins. Shown are (clockwise) OsNIP2;1, *M. marburgensis* AqpM, *E. coli* GlpF, SoPIP2;1 and HsAQP10. The fifth SF residue (G or T) is labeled in bold in classical aquaporins. The OsNIP2;1 channel is indicated with an asterisk. (E) Membrane plane slab surface view showing the cytoplasmic block of the channel caused by loop D (red). SF residues and the asparagines of the two NPA motifs are shown as sticks.

Inspection of the extracellular mouth of the channel reveals that the “GSGR” SF includes a fifth residue, Thr65 (Figs. 1C,D). In most other AQP and AQGP structures, the SF has only four residues due to the presence of an aromatic residue at the first position (*e.g*. **F**ISR in AqpM), which occludes the conserved glycine or threonine that corresponds to Thr65 in OsNIP2;1 (Fig. 1D). Thus, our structure suggests that plant Si transporters have a novel, 5-residue SF (TGSGR) that is substantially wider than the 4-residue SF of conventional AQPs (Fig. 1D). Intriguingly, a similar, but as yet unrecognised 5-residue SF (TGGIR) is present in human AQP10 (ref. 12), a known AQGP which was recently shown to transport Si at levels that may be physiologically relevant (13). However, OsNip2;1 transports glycerol very poorly (5), and we propose that this is due to the lack of a large hydrophobic SF residue (*e.g*. Trp and Phe in GlpF; Ile in AQP10), which is known to interact with the backbone of the amphipathic glycerol molecule (5). It is clear, at least from the static crystal structure, that the OsNIP2;1 SF is unique in having four oxygen atoms lining the channel, providing multiple hydrogen bond donor and acceptor groups for the four hydroxyl groups of the translocating Si molecule.

On the cytoplasmic side, the OsNIP2;1 channel is completely closed by loop D (residues ^185^ ATDTRA ^191^; Fig. 1E), suggesting protein-based regulation of transport activity, possibly via (de)phosphorylation. This is surprising, because Si is not known to be toxic, and in plants, excess Si would be deposited as silica bodies. Moreover, *lsi1* expression is downregulated upon continuous Si exposure (5), and no phosphorylation of OsNIP2;1 has yet been shown. On the other hand, loop D is conserved in NIPs, suggesting functional importance. Classical water-specific plant aquaporins such as PIP2;1 from spinach (SoPIP2;1) utilise phosphorylation-mediated loop D gating to protect the plant from drought stress and floods, conditions which require the channel to close (15). No evidence for phosphorylation was detected for yeast-expressed FL and truncated OsNIP2;1 by mass spectrometry analyses (Fig. S1) and inspection of the electron density, suggesting that channel opening might require phosphorylation. To test this notion and assess the significance of the closed OsNIP2;1 channel observed in the crystal, we performed equilibrium molecular dynamics (MD) simulations on the tetrameric assembly embedded in a POPC bilayer (Table S2). Strikingly, by taking the distance between Phe121 in loop B and R189 in loop D as a proxy for channel opening, three independent 0.5 μs simulations show a pronounced shift (up to ~15 Å for the Arg189 side chain) of loop D that opens the channel (Figs. 2A,B). Most (10 out of 12) monomers showed either a partial or complete channel opening (Fig. 2B and Fig. S3), suggesting the closed conformation of the channel was selected by the crystallisation process. HOLE profiles show that even the closed channels in the MD simulations are more open than the crystal structure (Fig. 2C). As expected from the fact that OsNIP2;1 also transports water (6), the channels fill with water on a picosecond time scale during system equilibration. Calculation of bidirectional water flux gives a reasonable correlation with the open/closed state of the channel. In OsNIP2;1_WATER1_ (Fig. 2B), for example, the tetrameric flux corresponds to 7.1 waters/ns, taken over the entire 500 ns of the simulation. However, the mostly open channel C (yellow) accounts for 2.8 waters/ns, and the mostly closed channel D (red) for 0.9 waters/ns. As a comparison, the glycerol-specific AQP7 (ref. 15), which also transports water very efficiently, yields a slightly higher flux of 12 waters/ns for the tetramer.

**Figure 2.**
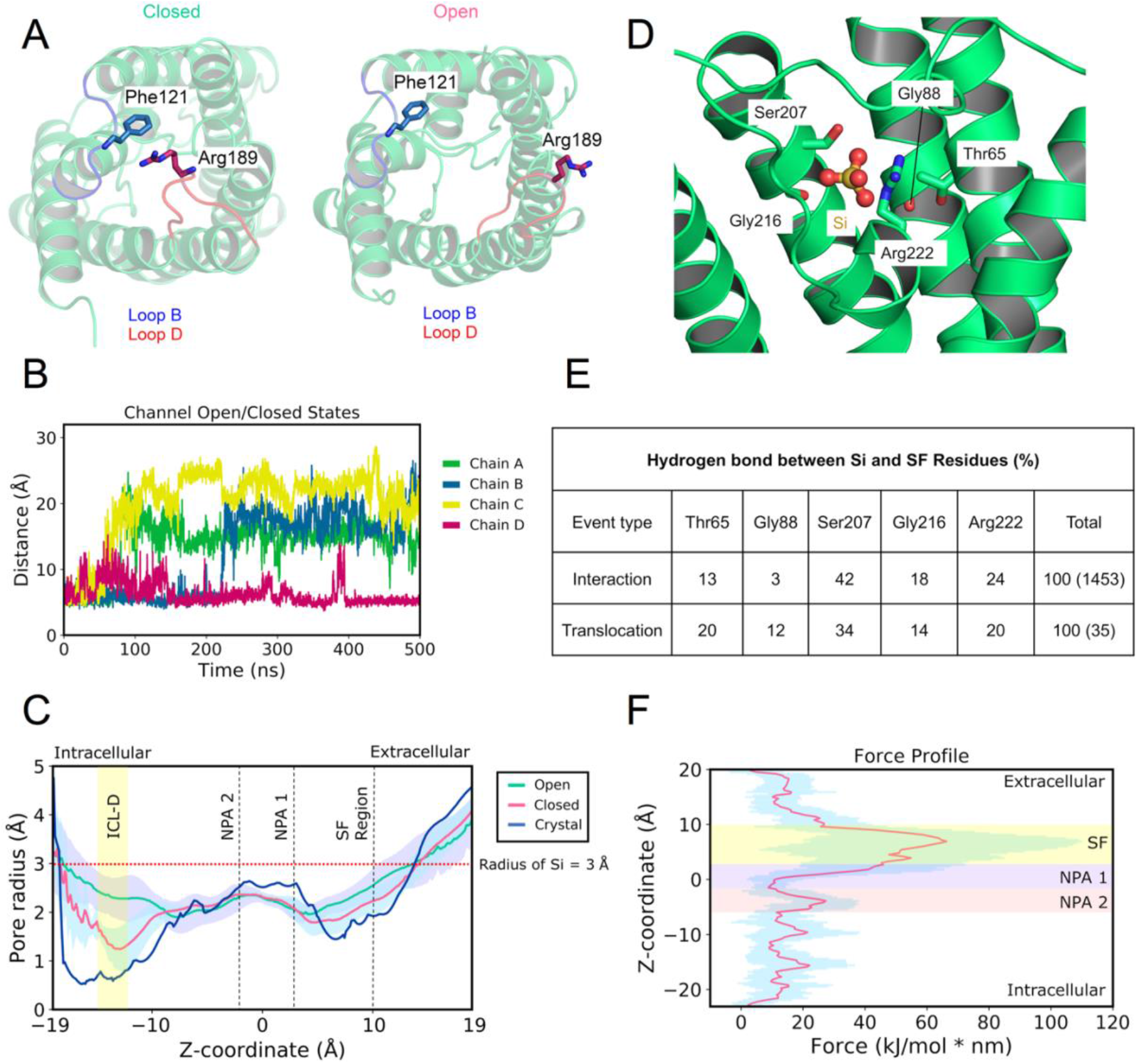
Molecular dynamics simulations of OsNIP2;1. (A) Comparison of the closed (left panel) and open channel structures. (B) Channel opening in the tetramer vs. simulation time. (C) HOLE profiles for the X-ray crystal structure and open and closed states from the MD simulations. The radius of the Si molecule is indicated for reference. (D) Simulation snapshot of a Si molecule in the SF. (E) Statistics for hydrogen bonds between Si and SF residues for all interaction events and translocations only. (F) Steered MD force profile for Si translocation. Average (red) value and standard deviations (blue) are shown for ten simulations.

We next generated a system with Si on both sides of the bilayer and performed unbiased MD simulations. Initial setups with 0.1 and 0.5 M Si did not give any translocation events in 250 ns simulations, and we therefore used 1 M Si. In total, 3 μs of simulations were performed in the presence of Si, yielding a total of three uptake events (defined as movement from the extracellular side into the space corresponding to the cytoplasm; Supplementary Movie). The translocating Si molecules form hydrogen bonds with all residues of the SF including Thr65 (Figs. 2D,E), but Ser207 appears to be especially important. Only a small fraction of interaction events at the SF result in translocation, suggesting that passage through the SF constitutes a thermodynamic or kinetic energetic barrier. Interestingly, the diameter of Si (6 +/- 0.5 Å) is larger than that of most of the channel, as determined via HOLE. This apparent discrepancy is due to the fact that HOLE calculates diameters based on the largest sphere that fits the channel. Given that both the channel and Si are non-spherical, this leaves enough space for permeation, but it does not allow co-permeation of Si hydration waters. As expected from its symmetry, the translocating Si molecule has no fixed orientation and rotates during its passage through the channel (Fig. S4).

In addition to three spontaneous uptake events, we also observed three export events, with Si moving from the cytoplasmic side to the extracellular side. In these cases, prior to entering the channel, the Si interacts extensively with residues located at the channel entrance, especially His106, Arg119 and Asp187 (Fig. S5). Interactions with the same residues are observed during uptake, after Si exits the channel proper and prior to diffusion away from the protein. Thus, although the sample size is small, the similar number of Si uptake and export events suggests that, in the absence of a gradient across the bilayer, Si transport would be bidirectional.

To increase the number of translocation events without the need for excessively long unbiased simulations, steered MD simulations were performed to provide further information about the location of translocation barriers. The SF region combined with the NPA1 motif (z ~ 10 to 0 Å) provides the largest barrier for the Si molecule, with a large deviation which indicates that the size of the barrier is dependent on the conformational arrangement of the amino acids in these regions. A second, smaller, barrier exists for the NPA2 domain (z ~ −2 to −6 Å), which the Si molecule seems to cross relatively easily in the simulations. Residence times for the Si in each of these regions were 16.0 ns ± 1.6 for the SF, 9.9 ns ± 2.5 for NPA1, and 8.4 ns ± 1.3 for NPA2 (Methods). These values correspond reasonably well to those obtained from the spontaneous permeation events (SF, 18.8 ns ± 13.5; NPA1, 8.3 ns ± 4.7; NPA2, 3.5 ns ± 2.7), and confirm that the SF presents the largest permeation barrier for Si translocation.

Silicon is a very important element for plants (1), and accumulating evidence suggests that it has health benefits for humans as well (2). Our results provide a platform that will lead to an improved understanding of Si uptake by plants that may be utilised for silicon biofortification of important crops. Within the wider context of metalloid transport, the study of NIP family members at the atomic level will generate insights into metalloid selectivity, that, together with *in vitro* and *in planta* studies, may enable rational manipulation of, *e.g*., boron levels in barley and arsenic accumulation in the rice grain.

## Author contributions

MD simulations were performed by CP, supervised by SK. JRB performed the mass spectrometry experiments. AB collected X-ray crystallography data. BvdB conceived the project, cloned, expressed and purified OsNIP2;1, and determined the crystal structures. The manuscript was written by BvdB with input from SK and CP.

## Acknowledgements

The authors thank Bastien Belzunces for calculation of the charges on silicic acid. The authors acknowledge the use of the IRIDIS High Performance Computing Facility, and associated support services at the University of Southampton and the use of the UK national supercomputer, ARCHER granted via the UK High-End Computing Consortium for Biomolecular Simulation, HECBioSim (http://hecbiosim.ac.uk), supported by EPSRC (grant no. EP/R029407/1), in the completion of this work. SK is supported by the same grant. Research in C.V.R.’s laboratory is supported by a Medical Research Council program grant (MR/N020413/1).

## Methods

### OsNIP2;1 expression and purification

The gene sequence encoding for full-length *osnip2;1* was obtained by gene synthesis (Eurofins genomics) and optimised for expression in *Saccharomyces cerevisiae*. Three putative N-terminal N-glycosylation sites were removed via replacement with glutamines (N4Q/N13Q/N26Q). The resulting gene, encoding a hexa-histidine sequence at the C-terminus for purification, was cloned via BamHI and XhoI restriction sites into the 83ν vector (17) digested with the same enzymes. The plasmid was moved into the yeast W303 Δ*pep4* expression strain via the lithium acetate method. Transformants were selected on SCD -His plates (ForMedium), incubated at 30 °C.

For expression, cells were grown in shaker flasks at 30 °C for ~20-24 hrs in synthetic minimal medium lacking histidine and with 1% (w/v) glucose to a typical OD600 of 6-8. Cells were subsequently spun down for 15 mins at 4200 rpm and resuspended in YP medium containing 1.5% (w/v) galactose, followed by another 16-20 hrs growth at 30 °C/225 rpm, and harvested by centrifugation for 20 mins at 4200 rpm. Final OD_600_ values typically reached 18-20. Cells were resuspended in TSB buffer (20 mM Tris, 300 mM naCl, pH 8) in the presence of 5 mM EDTA and lysed by 1-2 passes through a cell disrupter operated at 35-37 kpsi (TS-Series 0.75 kW; Constant Systems). Membranes were collected from the suspension by centrifugation at 200,000 x g for 90 mins (45Ti rotor; Beckmann). Membrane protein extraction was performed by homogenisation in TSB with a 1:1 (w/w) mixture of dodecyl-β-D-maltoside and decyl-β-D-maltoside (DDM/DM) followed by stirring at 4 °C for 1 hr or overnight. Depending on the amount of processed cells, one or two protease inhibitor tablets were added at this stage (Complete EDTA-free protease cocktail; Sigma). Typically, 1 g (1% w/v) of total detergent was used for membranes from 2 liters of cells. The membrane extract was centrifuged for 35 mins at 200,000 x g and the supernatant loaded onto a 10 ml Nickel column (Chelating Sepharose; GE Healthcare) equilibrated in TSB with 0.2% DDM pH 8. The column was washed with 15 column volumes buffer containing 30 mM imidazole and eluted in 3 column volumes with 200 mM imidazole. The protein was purified to homogeneity by gel filtration chromatography in 10 mM Hepes, 100 mM NaCl, 0.05% DDM pH 7-7.5. For polishing and detergent exchange, a second gel filtration column was performed using various detergents. The final yield of OsNIP2;1 was about 1 mg per 4 liters of culture. Proteins were concentrated to ~10-15 mg/ml using 100 kD cutoff centrifugal devices (Millipore), flash-frozen and stored at −80 °C.

### Crystallisation and structure determination

Crystallisation screening trials by sitting drop vapor diffusion were set up at 4 °C and 20 °C using in-house screens and the MemGold, MemGold2, MemChannel and MemTrans screens (Molecular Dimensions) with a Mosquito crystallisation robot (TTP Labtech). Crystals were harvested directly from the initial trials or optimised by hanging drop vapor diffusion using larger drops (typically 2-3 μl total volume). Crystals diffracting beyond 5 Å were obtained only with 0.05% decyl-maltose neopentyl glycol (DMng), with the best crystal showing useable diffraction to just beyond 4 Å (C2 space group). However, the diffraction was anisotropic, and molecular replacement solutions could not be obtained. To improve diffraction, limited proteolysis trials were performed at 4 °C with trypsin and chymotrypsin, using a 100-fold excess (w/w) of OsNIP2;1 over protease. The digestions with trypsin showed removal of ~5-10 kD from the protein based on SDS-PAGE. Following a large-scale digest (~ 5 mg), the truncated protein was subjected to SEC in DMng as described above, with the addition of 10% glycerol to the buffer. Native mass spectrometry showed a molecular mass for the monomer of 23960 Da, indicating the likely removal of residues 1-37 from the N-terminus and residues 265-304 from the C-terminus (predicted molecular mass 23964.1 Da). After initial screening as described above, diffracting, block-shaped crystals with various morphologies in space group P1 were obtained following optimisation of the MemGold H11 hit condition (1 mM CdCl_2_, 30 mM MgCl_2_.6H_2_O, 0.1 m MES pH6.5, 30% PEG400), by slightly varying the PEG 400 concentration (28-32% w/v). Due to severe anisotropy in most crystals, a number of crystals had to be screened in order to obtain a moderately anisotropic dataset that allowed successful structure solution. Datasets (360 degrees) were collected at beamline I24 at the Diamond Light Source and were autoprocessed via XDS within Autoproc (18) and STARANISO (19). Useful phases were obtained via molecular replacement (MR) with Phaser (20), using as search model the tetramer of the AqpM aquaporin from *Methanothermobacter marburgensis* (PDB ID: 2EVU), which has 34% sequence identity to OsNIP2;1. The asymmetric unit (AU) contains two OsNIP2;1 tetramers, corresponding to a solvent content of ~65% (Matthews volume 3.5). The two tetramers within the AU pack via their cytoplasmic faces, in part via bridging cadmium ions present in the crystallisation mixture. However, it is not clear whether the cadmium ions play a critical role in lattice formation, given that the C2 crystals for the full-length protein, obtained in the absence of cadmium, pack in a very similar manner. The initial model was improved via iterative cycles of Autobuilding within Phenix (21), manual building within Coot (22), and refinement via Phenix. The data for refinement were cut off at 3.0 Å, since higher resolution cutoffs led to unstable refinements with high clash scores and many rotamer and Ramachandran outliers (> 3%). Structure validation was carried out with MolProbity (23). The data collection and refinement statistics are summarized in Table S1.

### Native mass spectrometry

Prior to MS analysis, the protein sample was buffer exchanged into 200 mM ammonium acetate pH 8.0 and 0.05% (w/v) LDAO, using a Biospin-6 (BioRad) column and introduced directly into the mass spectrometer using gold-coated capillary needles (prepared in-house). Data were collected on a Q-Exactive UHMR mass spectrometer (ThermoFisher). The instrument parameters were as follows: capillary voltage 1.2 kV, quadrupole selection from 1,000 to 20,000 m/z range, S-lens RF 100%, collisional activation in the HCD cell 100 V, trapping gas pressure setting kept at 7.5, temperature 200 °C, resolution of the instrument 12500. The noise level was set at 3 rather than the default value of 4.64. No in-source dissociation was applied. Data were analysed using Xcalibur 4.2 (Thermo Scientific).

### Proteomics

For protein identification, proteins were digested with both trypsin and chymotrypsin and the resultant peptides were loaded onto a reverse phase C18 trap column (Acclaim PepMap 100, 75 μm x 2 cm, nano viper, C18, 3 μm, 100 Å, ThermoFisher, Waltham, MA, U.S.A) using an Ultimate 3000 and washed with 50 μL of solvent A at 10 μl/min. The desalted peptides were then separated using a 15 cm pre-packed reverse phase analytical column (Acclaim PepMap 100, 75μm x 15 cm, C18, 3 μm, 100 Å, ThermoFisher, Waltham, MA, U.S.A) using a 45 min linear gradient from 5% to 40% solvent C (80% acetonitrile, 20% water, 0.1% formic acid) at a flow rate of 300 nl/min. The separated peptides were electrosprayed into an Orbitrap Eclipse Tribrid mass spectrometry system in the positive ion mode using data-dependent acquisition with a 3 s cycle time. Precursors and products were detected in the Orbitrap analyzer at a resolving power of 60,000 and 30,000 (@ m/z 200), respectively. Precursor signals with an intensity >1.0 × 10^−4^ and charge state between 2-7 were isolated with the quadrupole using a 0.7 m/z isolation window (0.5 m/z offset) and subjected to MS/MS fragmentation using higher-energy collision induced dissociation (30% relative fragmentation energy). MS/MS scans were collected at an AGC setting of 1.0 x 10^4^ or a maximum fill time of 100 ms and precursors within 10 ppm were dynamically excluded for 30 s. Data were searched against the Oryza sativa subsp. japonica (Rice) proteome using ProteinProspector (v6.2.2) with the following search parameters: trypsin/chymotrypsin digestion; fixed modification was set to carbamidomethyl (C); variable modifications set as oxidation (M), acetylated protein N terminus, and phosphorylation (STY).

### Molecular Dynamics Simulations

Systems were constructed by submitting the crystallographic structure of OsNIP2;1 to the CHARMM-GUI server (24,25) and using the Membrane Builder feature (26) (Wu et al., 2014). The aquaporin was inserted in a membrane composed of 1-palmitoyl-2-oleoyl-sn-glycero-3-phosphocholine (POPC) phospholipids and with explicit water solvation. A number of counterions were added to the simulation to neutralize charges, with an extra salt concentration of 0.15 M of potassium chloride ions for all simulations. Concentrations of 0.1, 0.5, and 1 M of Si were tested, aiming to increase the number of translocation events observed (uptake/export). Si molecules were randomly inserted in the simulation box by making use of the *gmx insert-molecules* tool. Si molecule topology was constructed using pre-existing silicate parameters (27) from the CHARMM36m force field (28). Atomic partial charges for Si were obtained using quantum mechanics (QM) calculations of Hirshfeld charges (29) in Gaussian16 software (30). For this, an optimization of the system was performed at the MP2 level (31–35) with a 6-311G** basis set (36,37).

Molecular dynamics simulations were performed using the GROMACS simulation suite (version 2020.4) (38) along with CHARMM36m force field (28,39) and TIP3P water model (40). Initially, equilibration simulations were run employing NVT and NPT ensembles with position restraints in the protein and phosphate atoms of the phospholipids, lasting for 1 and 50 ns, respectively. Subsequentially, production simulations in NPT ensemble ran for either 500 ns or 1 microsecond. Simulations were performed at both 310 K and 330K (to enhance the occurrence of Si translocation events), and the velocity rescale thermostat (41) with a coupling constant of τ = 0.1 was applied to keep a constant temperature. The Parrinello-Rahman barostat (42) with a time constant of 2 ps was used to maintain pressure semi-isotropically at 1 atm. Long-range electrostatics were treated by the particle mesh Ewald method (43). Covalent bonds were constrained by the LINCS algorithm (44,45), allowing an integration step of 2 fs. Values for long-range electrostatics and van der Waals cut-offs were set to 1.2 nm. Starting velocities were modified at the beginning of each different replicate to improve conformational sampling. Ten steered MD simulations were performed for 40 ns to estimate the force required for one Si molecule to translocate through the channel. To achieve this, a harmonic spring constant with a force of 800 kJ mol^−1^ nm^−2^ was attached to the Si atom, which was then pulled through the z-axis at a constant velocity of 0.1 nm ns^-1^. Different initial starting velocities were employed for each of the ten independent runs. Final forces were computed and an average force profile and standard deviation were obtained for the pulling coordinate.

Molecules were manipulated, visualized, and analysed utilizing VMD (46) and Pymol (47) software. The HOLE software (48) was utilized for the calculation of pore radius analyses. Distance and angle cut-offs to count hydrogen bonds between atoms were 3.5 Å and 20º, respectively. A translocation event was counted once a Si molecule moved from one side of the simulation box, through the inside of the protein channel, and exited on the opposite side. Residence time analysis was performed by calculating the minimum distance between Si and the atoms of each domain. The molecule was considered to be interacting with the domain if the distance was less than 4 Å. Once this distance was inside the cut-off, the number of subsequent frames that it remained below the cut-off was calculated and converted to the corresponding amount of time (in ns). The HOLE profiles from the open and closed states were obtained by dividing the trajectories into 0-200 and 300-500 ns portions. A total of 12 portions were used (6 for each state) in the analysis, using data from different channels, followed by calculation of the average profile and standard deviation. For the closed state, portions were chosen if distance values between R189 and F125 were below 10 Å: chain B (OsNIP2;1_WATER1_), chains A, B, and C (OsNIP2;1_WATER2_), chain A (OsNIP2;1_WATER3_). For the open state, portions were chosen when distance values were above 10 Å: chains A, B, and C (OsNIP2;1_WATER1_), chain B (OsNIP2;1_WATER2_), chain A and C (OsNIP2;1_WATER3_).

### Accession codes

Coordinates and structure factors have been deposited in the Protein Data Bank with accession code 7NL4 [http://doi.org/10.2210/pdb7NL4/pdb].

---

**Figure S1.**
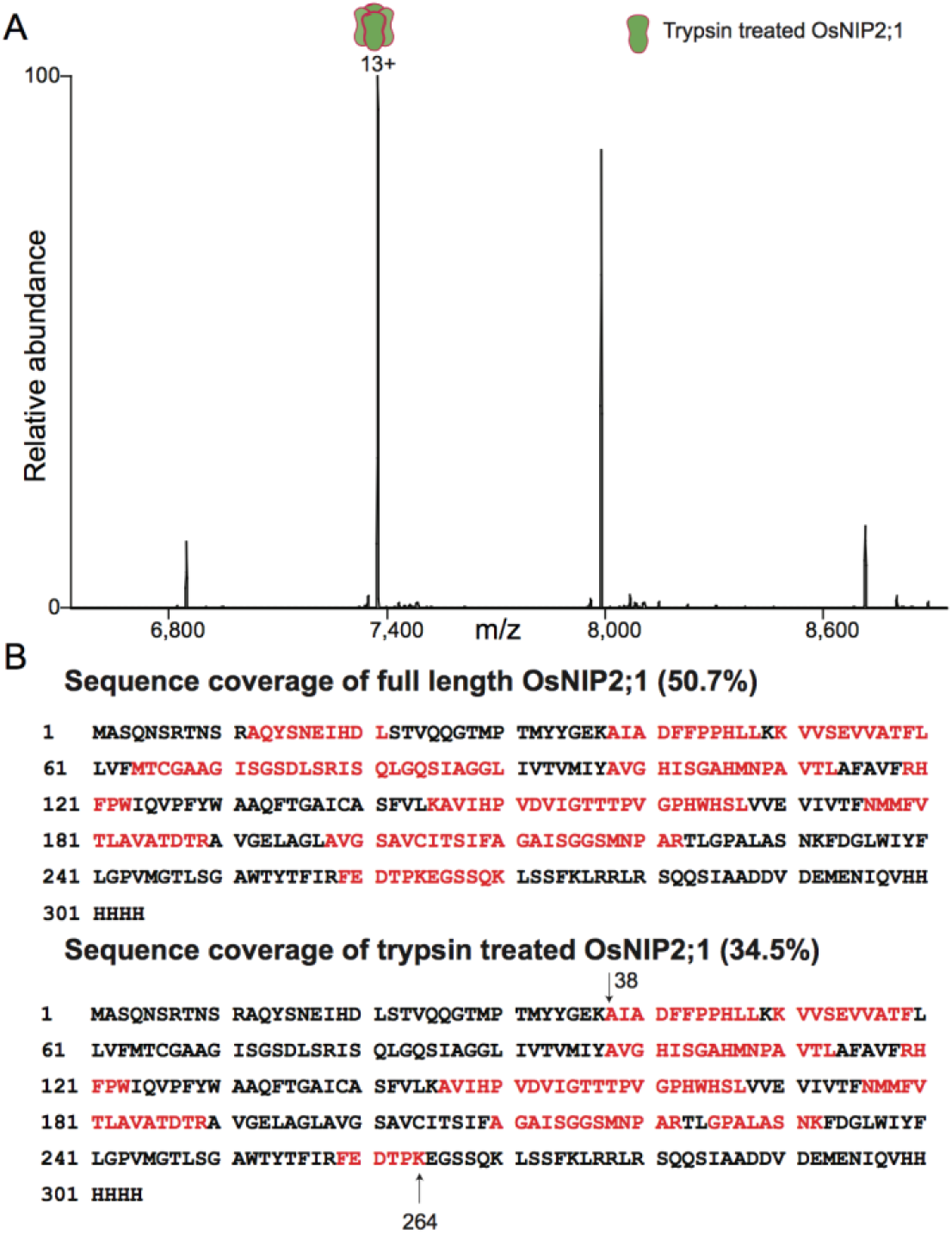
Mass spectrometry analyses of OsNIP2;1. (A) Native mass spectrum of trypsin-treated OsNIP2;1 in LDAO reveals a charge state series consistent with a tetrameric assembly of OsNIP2;1. The observed mass for the tetramer is 95856±1 Da and the respective collision-induced dissociated monomer mass of 23,962 Da is close to the theoretical value for the fragment corresponding to residues 38-264 (23,964.1 Da). (B) Observed peptides (red) after digestion of full-length (top) and truncated OsNIP2;1 (bottom) with trypsin and chymotrypsin identified by proteomics. Calculated mass from these observed peptides (38-264) in the case of truncated protein aligns quite well with the mass observed in native mass spectrometry data in (A).

**Figure S2.**
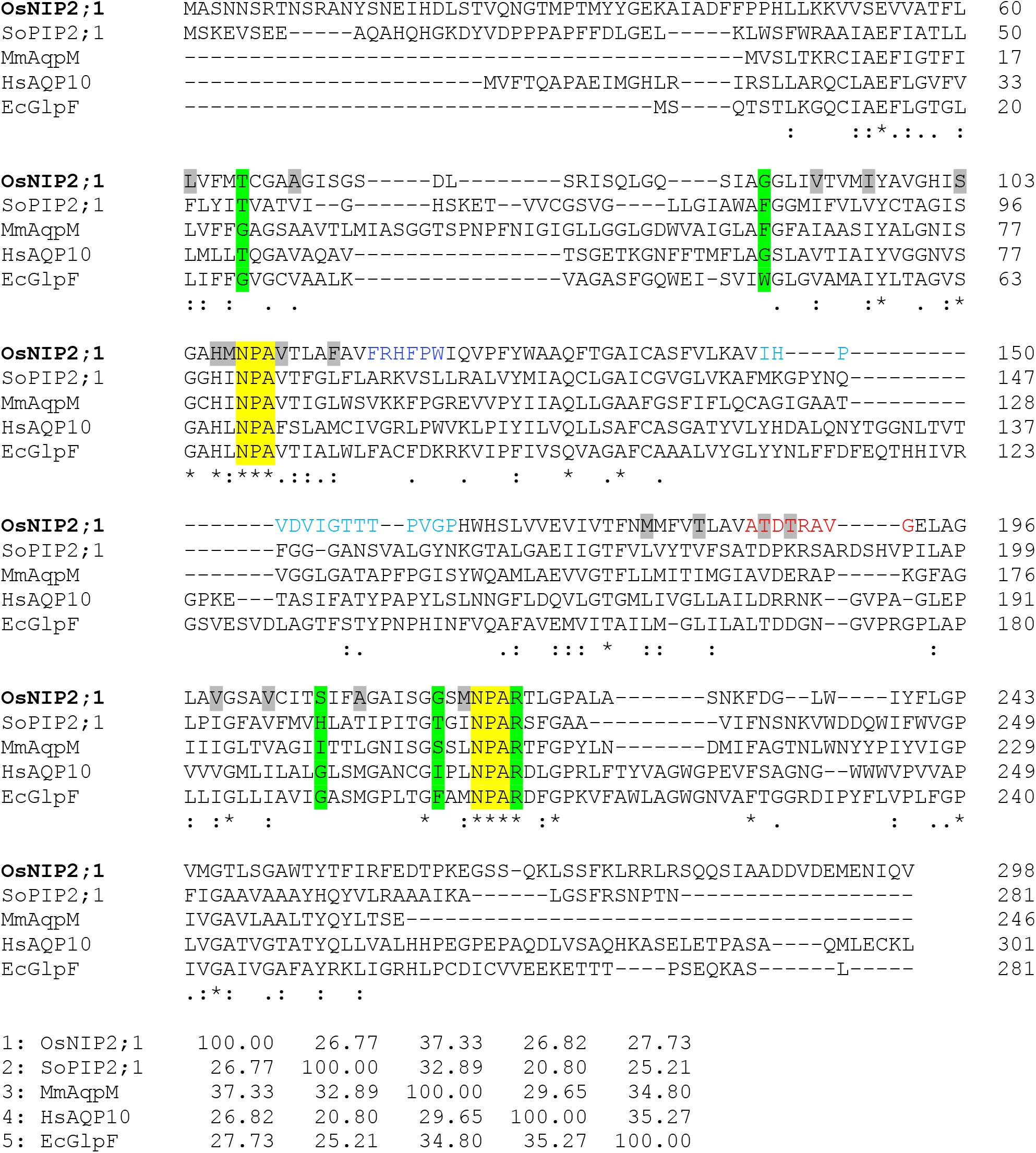
Clustal omega sequence alignment of OsNIP2;1 with other aquaporins. The following sequences were analysed: *Oryza sativa* NIP2;1 (rice), *Spinaca oleracea* PIP2;1 (spinach), *Methanothermobacter marburgensis* AqpM, human AQP10 and *E. coli* GlpF. Selectivity filter residues are highlighted in green, and the NPA motifs in yellow. Residues lining the OsNIP2;1 channel are highlighted grey. OsNIP2;1 residues of loops B, C and D are shown in dark blue, cyan and red, respectively. The bottom panel shows the identity matrix for the five aquaporins.

**Figure S3.**
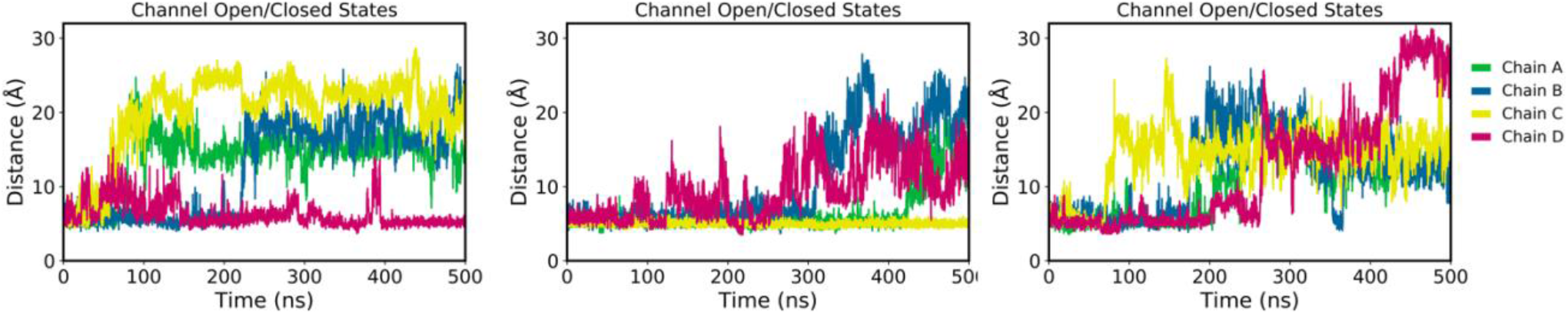
Channel opening during MD simulations. Each plot shows the distance between the R189 guanidium group and the Cγ atom from F125 side-chain, highlighting that only two channels did not open at all when considering the three simulations together. From left to right, Water_R1_, Water_R2_ and Water_R3_ simulations are shown, respectively.

**Figure S4.**
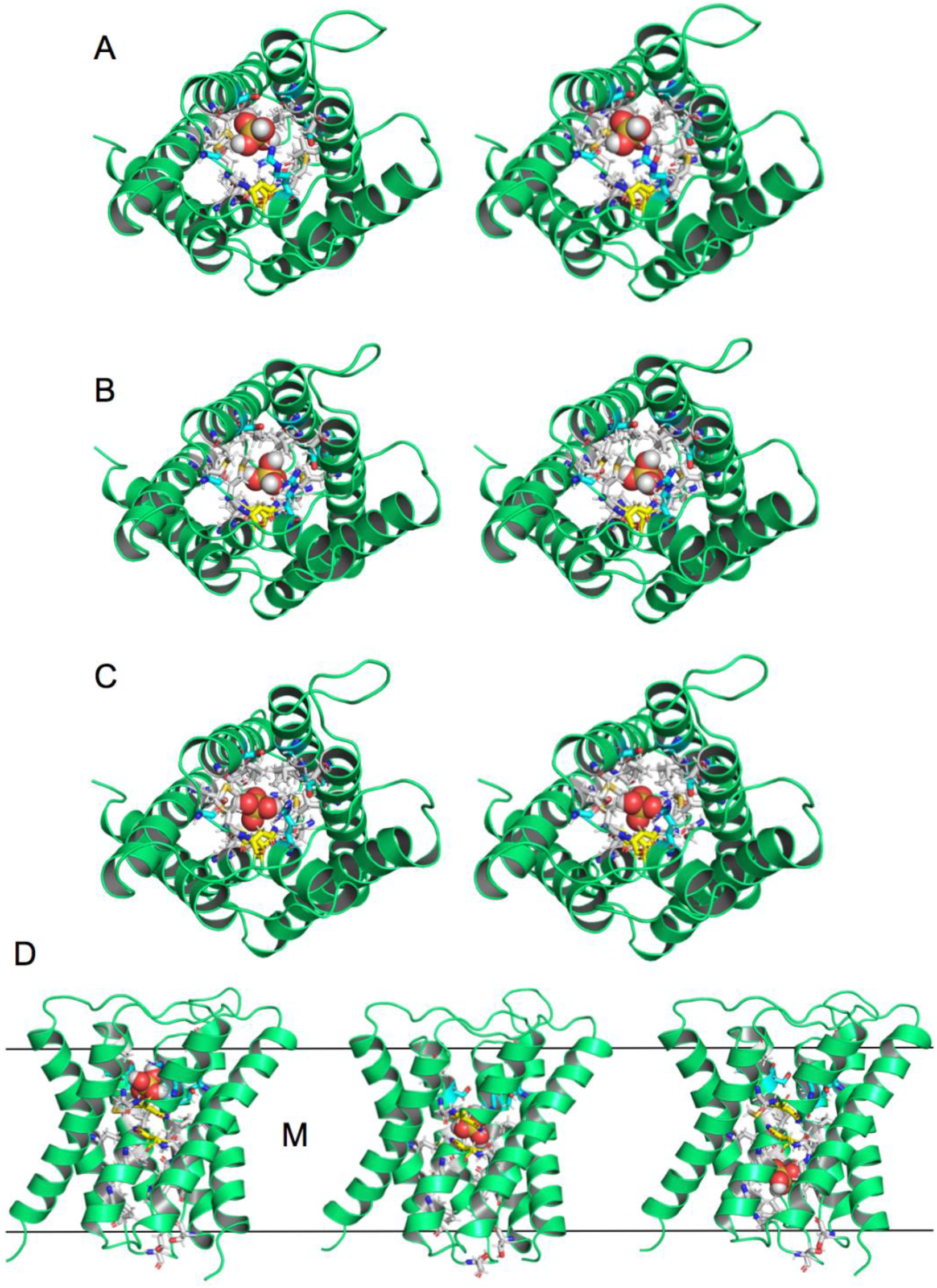
Si translocation through OsNIP2;1. (A-C) Stereo view snapshots viewed from the extracellular side of Si interacting with the SF (A), NPS motifs (B) and with H106 in the exit site (C). (D) Views from the plane of the membrane, with the membrane boundaries indicated. SF residues are coloured cyan, N108 and N219 are yellow, and other channel-lining residues are grey. The Si molecule is shown as a space-filling model.

**Figure S5.**
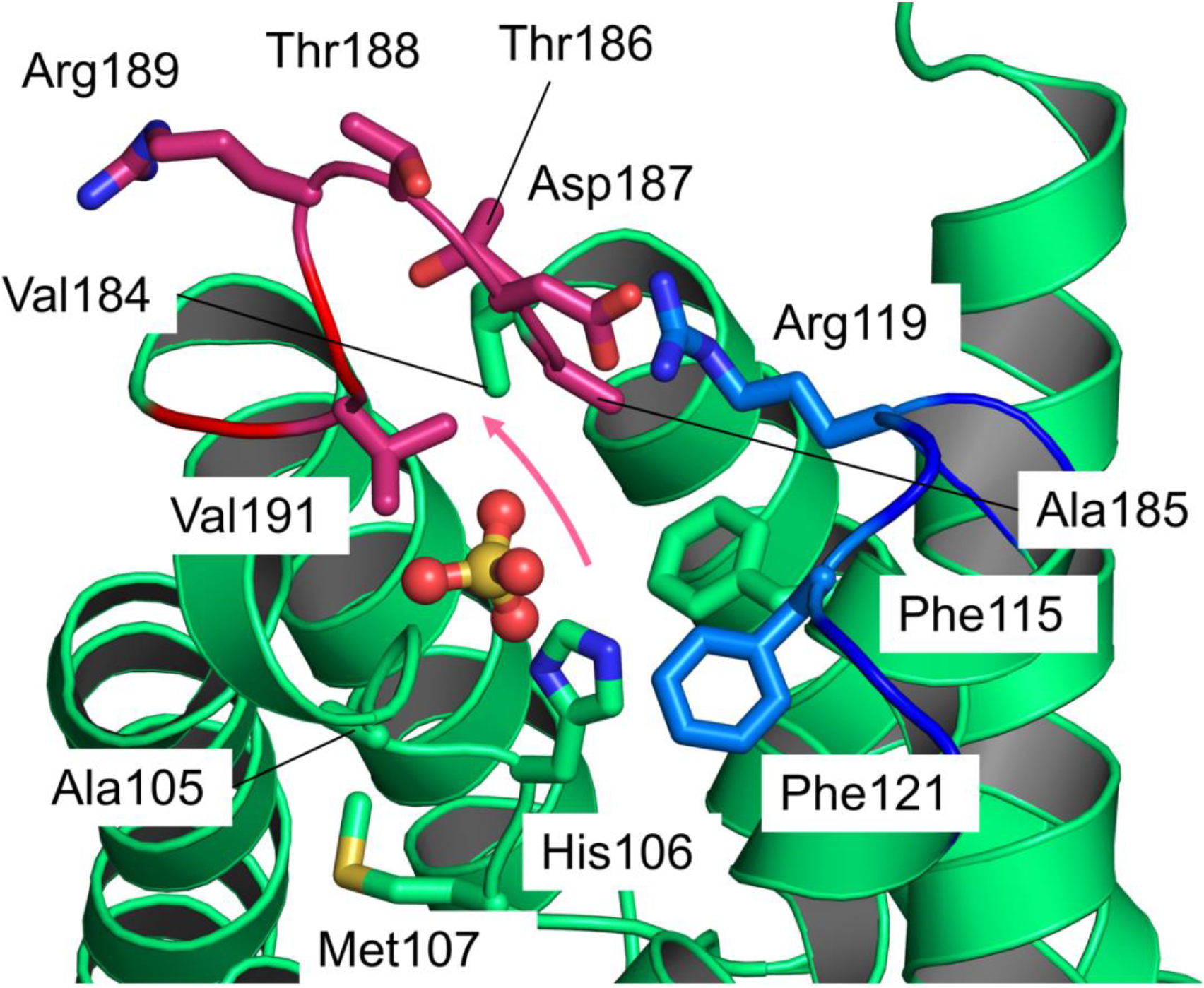
The cytosolic binding site of OsNIP2;1. Zoomed-in image of the cytosolic binding site of OsNIP2;1, showing important residues in this region of the protein. The protein is shown as a green cartoon. Loop B is coloured dark blue, while Loop D is coloured red. Relevant residues are shown as stick models. The Si molecule is shown as a ball-and-stick representation, with a pink arrow indicating the movement performed by the molecule to exit the binding site.

**Table S1.**
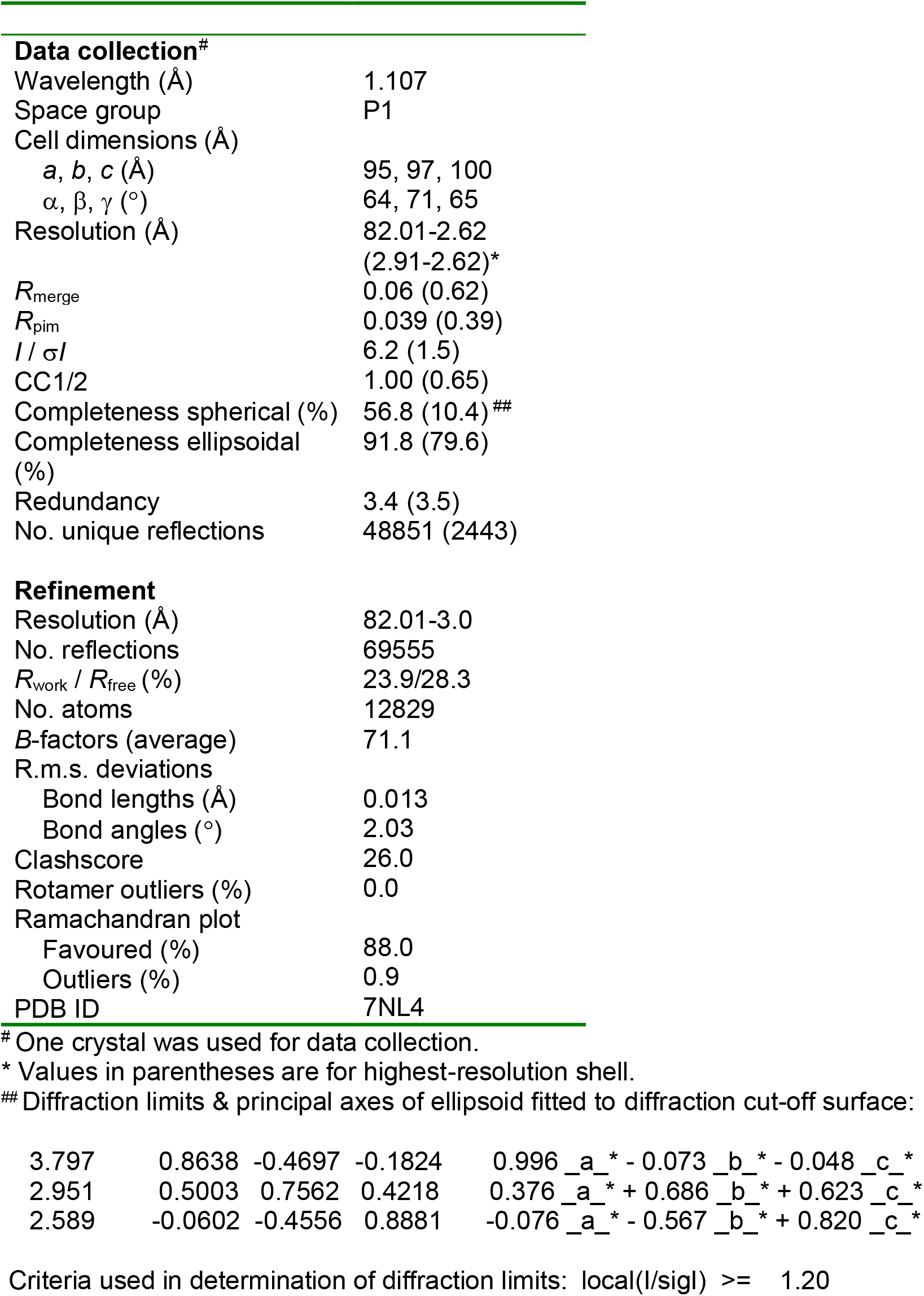
X-ray crystallographic data collection and refinement statistics for OsNIP2;1

**Table S2.** Summary of molecular dynamics simulations.

| System | Length | Si concentration | Si initial distribution |
| --- | --- | --- | --- |
| OsNIP2;1<sub>WATER</sub> | 3 x 0.5 μs | - | - |
| OsNIP2;1<sub>LOWSi</sub> | 1 x 0.25 μs | 0.1 M | No Si bound to OsNIP2;1 |
| OsNIP2;1<sub>MEDIUMSi</sub> | 1 x 0.25 μs | 0.5 M | No Si bound to OsNIP2;1 |
| OsNIP2;1<sub>HIGHSi</sub> | 2 x 1 μs | 1.0 M | No Si bound to OsNIP2;1 |
| OsNIP2;1<sub>SFSi</sub> | 2 x 0.5 μs | 1.0 M | One Si at SF* |

\* In both of these simulations Si moved rapidly out of the SF into the extracellular bulk solution. The initial position of the Si, where it interacts with the SF, was not included in the hydrogen bond statistics calculation of Fig. 2e.

Supporting Movie. Si translocation by OsNIP2;1. Unbiased molecular simulation trajectory showing a translocation event of a Si molecule through one aquaporin monomer. Colour scheme as above, hydrogens are not shown.

